# Improved production and expanded application of CVS-N2c-ΔG virus for retrograde tracing

**DOI:** 10.1101/2022.01.22.477330

**Authors:** Kunzhang Lin, Lei Li, Wenyu Ma, Xin Yang, Zengpeng Han, Nengsong Luo, Fuqiang Xu

## Abstract

Neurotropic virus tracers, particularly those with low toxicity and high efficient tracing, are powerful tools for structural and functional dissections of neural circuits. The retrograde trans-mono-synaptic technology based on rabies virus CVS-N2c strain has reduced cytotoxicity and enhanced efficiency, attains long-term gene manipulation for functional studies, but suffers from difficult preparation and low yield. To overcome these shortcomings, an improved production system was established for rapid rescue and preparation of CVS-N2c-ΔG virus, CVS-N2c-ΔG with the same titer as SAD-B19-ΔG can be prepared within a short time. Meanwhile, we found that N2cG coated CVS-N2c-ΔG allows efficient retrograde access to projection neurons, and further expand its application in VTA/SNc to DLS pathway that unaddressed by rAAV9-Retro, and the efficiency is 6 folds higher than that of rAAV9-Retro. Then the trans-synaptic efficiency of CVS-N2c-ΔG virus was evaluated. Results showed that the trans-mono-synaptic efficiency of oG-mediated CVS-N2c-ΔG was 2-3 folds higher than that of oG-mediated SAD-B19-ΔG, but there was no difference between oG-mediated and N2cG-mediated CVS-N2c-ΔG system. In addition, codon modified N2cG (optiG) did not increase the efficiency of CVS-N2c-ΔG tracing. Finally, we found that the CVS-N2c-ΔG produced by the improved method can be used for monitoring neural activity of projection neurons, and the time window can be maintained for 3 weeks, and it can also express sufficient recombinases for efficient transgene recombination. That is, the virus produced by the improved production system does not affect its own function, paving the way for its further optimization, popularization and application in structural and functional studies of neural circuits.

## Introduction

Analyzing the connection of brain neural networks, including input and output neural pathways, is the basis to reveal the principle of the brain function and elucidate the mechanism of brain diseases [1]. Structural and functional studies of brain connection require anterograde and retrograde viral tracers [2–5]. The retrograde trans-mono-synaptic technology based on rabies virus SAD-B19 strain can label the input network of specific types of neurons [6, 7], which has been used to solve a large number of neuroscience problems and has been widely popularized and applied in the field of neuroscience [8, 9]. However, this system enables trans-mono-synaptic retrograde labeling only a fraction of upstream neurons, which may lead to the neglect of related input network connections [10]. Moreover, the high cytotoxicity makes it not conducive to carrying functional genes for neural activity detection and functional manipulation for a long time. The chimeric glycoprotein (oG) obtained by codon optimization can increase the trans-mono-synaptic efficiency of SAD-B 19 up to 20 folds [11]. Since then, many laboratories have used adeno-associated virus expressing oG as an auxiliary vector to track the upstream input of neural network [12–14]. The self-inactivating rabies virus (SiR) developed by rapidly degrading the N protein of SAD-B19-ΔG that related to virus replication, can carry Cre or Flpo recombinase combined with adeno-associated virus expressing functional probe for the study of functional network [15]. However, it also has some defects, such as complex preparation process, weak self-expression and unable to express sufficient functional probes. Chatterjee et al reported non-toxic SAD-B19 (SAD-B19-ΔGL) with double deletion of G and L proteins, however, it has only very weak expression ability, and can only express recombinase to magnify gene expression (such as AAV virus or transgenic animals equipped with recombinase-dependent functional probes expression) for the activity monitoring and genetic manipulation of functional networks [16]. Moreover, due to the large size of L gene, it is necessary to adopt appropriate helper virus vector strategy or make transgenic animals expressing L to realize trans-mono-synaptic tracing [16]. Reardon et al. established a retrograde trans-mono-synaptic system modified by rabies virus CVS-N2c strain. Through reverse compensation of its own glycoprotein N2cG, CVS-N2c-ΔG exhibits significantly enhanced retrograde trans-synaptic ability and further reduced toxicity to neurons compared with SAD-B19-ΔG vaccine strain [10]. CVS-N2c-ΔG can express functional probes such as calcium-sensitive probes and optogenetic probes for the analysis of functional network, therefore, it has more advantages in neural circuit tracing, but also has the disadvantages such as difficult preparation and low yield. Therefore, it has not been popularized in the following application.

Here, to overcome these limitations, we developed a new production system for rapid rescue and preparation of CVS-N2c-ΔG virus. The CVS-N2c-ΔG virus permits efficient retrograde labeling of projection neurons unaddressed by rAAV9-Retro, and maintains excellent performances for retrograde trans-mono-synaptic targeting, functional monitoring and transgene recombination.

## Results

### Optimized preparation method for CVS-N2c-ΔG viruses

CVS-N2c-ΔG exhibits significantly enhanced retrograde trans-synaptic ability and further reduced toxicity to neurons compared with SAD-B19-ΔG vaccine strain, and it can express functional probes such as calcium-sensitive probes and optogenetic probes for the analysis of functional network [10], but it also has the disadvantages of very prolonged preparation process and low yield. Therefore, it is necessary to establish a highly efficient preparation method of CVS-N2c-ΔG viruses. Here we provide a new preparation protocol involving three cell lines as shown in Fig. 1A: 1) “B7GG” cell line [17], based on BHK-21 cells stably expressing the T7 RNA polymerase, the nuclear-localized EGFP and the SAD-B19 glycoprotein (B19G), used for the rapid rescue and amplification of CVS-N2c-ΔG viruses; 2) “BHK-N2cG” cell line (Fig. S1A), based on BHK-21 cells stably expressing the CVS-N2c glycoprotein (N2cG), along with the nuclear-localized EGFP, used for the amplification of N2cG coated CVS-N2c-ΔG viruses; 3) “BHK-EnvARVG” cell line (Fig. S1B), based on BHK-21 cells stably expressing the EnvA and B19G chimeric glycoprotein (EnvARVG) [6], along with the nuclear-localized EGFP, used for the amplification of EnvARVG coated CVS-N2c-ΔG viruses. We found that the CVS-N2c-ΔG virus can be quickly rescued (Fig. 1B). To evaluate the production performance of these cell lines, we compared the production efficiency of CVS-N2c-ΔG and SAD-B19-ΔG viruses in these cells, by means of supernatant titer detection. In B7GG cells, no significant difference was observed in supernatant titer between CVS-N2c-ΔG and SAD-B19-ΔG viruses (Fig. 1C, SAD-B19: 1.04 ± 0.26 x 10^6^ IU/ml; CVS-N2c: 0.88 ± 0.30 x 10^6^ IU/ml; P = 0.6978); In BHK-N2cG cells, the supernatant titer of CVS-N2c-ΔG was lower than that produced in B7GG cell line (Fig. 1D, B19G: 0.88 ± 0.30 x 10^6^ IU/ml; N2cG: 1.04 ± 0.26 x 10^5^ IU/ml; P = 0.0338). In addition, the SAD-B19-ΔG virus can also be amplified in this cell line, and the supernatant titer was also significantly lower than that produced in B7GG cell line (Fig. 1E, B19G: 1.04 ± 0.26 x 10^6^ IU/ml; N2cG: 1.24 ± 0.25 x 10^5^ IU/ml; P = 0.0077). However, there was no significant difference in supernatant titer between CVS-N2c-ΔG and SAD-B19-ΔG produced in the same BHK-N2cG cell line (Fig. 1F, SAD-B19: 1.24 ± 0.25 x 10^5^ IU/ml; CVS-N2c: 1.04 ± 0.26 x 10^5^ IU/ml; P =0.5917).

**Fig. 1.**
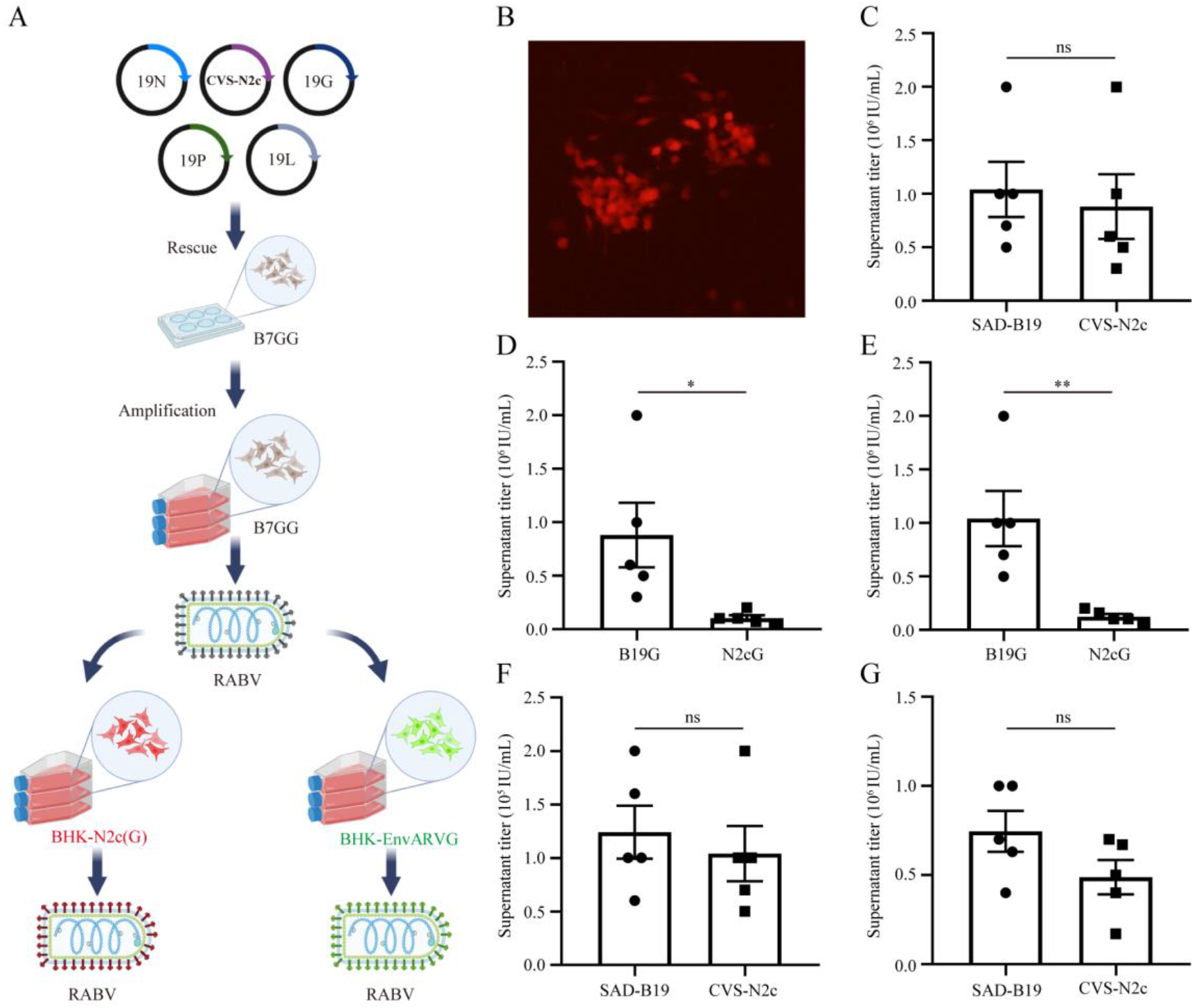
Optimized method for rapid rescue and preparation of CVS-N2c-ΔG virus. (A) Schematic diagram of virus production process using three different cell lines. “B7GG” cell line for the rapid rescue and amplification of CVS-N2c-ΔG viruses; “BHK-N2cG” cell line for the amplification of N2cG coated CVS-N2c-ΔG viruses; “BHK-EnvARVG” cell line for the amplification of EnvARVG coated CVS-N2c-ΔG viruses. (B) Fluorescent signals (red) of CVS-N2c-ΔG virus on the 5th day of rescue in B7GG cells. (C) Comparison of supernatant titers between CVS-N2c-ΔG and SAD-B19-ΔG viruses produced in B7GG. (D) Comparison of CVS-N2c-ΔG viral supernatant titers produced in B7GG and BHK-N2cG. (E) Comparison of SAD-B19-ΔG viral supernatant titers produced in B7GG and BHK-N2cG. (F) Comparison of supernatant titers between CVS-N2c-ΔG and SAD-B19-ΔG viruses produced in BHK-N2cG. (G) Comparison of supernatant titers between CVS-N2c-ΔG and SAD-B19-ΔG viruses produced in BHK-EnvARVG. Statistical values are indicated as mean ± SEM. Significant differences are expressed by the p value. *P<0.05, **P<0.01, ***P<0.001, ns, no significant difference.

It was previously reported that Neuro2A-EnvARVG was inefficient in producing CVS-N2c-ΔG virus [10], which may be because EnvARVG itself was not suitable for CVS-N2C-ΔG virus packaging, and may also be due to the low viability of Neuro2A cells. To verify whether BHK-EnvARVG cell line can efficiently produce high-titer EnvARVG coated CVS-N2C-ΔG, we used high-titer CVS-N2C-ΔG pseudotyped with B19G (produced by B7GG cell line) to infect BHK-EnvARVG cells. As shown in Fig. 1F, the supernatant titer was measured and compared with EnvARVG pseudotyped SAD-B19-ΔG. No significant difference were detected in supernatant titer between CVS-N2c-ΔG and SAD-B19-ΔG produced in the same BHK-EnvARVG cell line (Fig. 1G, SAD-B19: 7.46 ± 1.15 x 10^5^ IU/ml; CVS-N2c: 4.88 ± 0.97 x 10^5^ IU/ml; P = 0.1242), indicating that BHK-EnvARVG cell line can efficiently prepare high-titer EnvARVG pseudotyped CVS-N2C-ΔG virus.

These results showed that the optimized preparation method we designed can be used to produce high-quality and high-titer EnvARVG coated CVS-N2c-ΔG viruses for retrograde trans-mono-synaptic labeling.

### Retrograde access to projection neurons with CVS-N2c-ΔG virus

The retrograde labeling of viral tracers can be used to analyze the upstream neural networks projected to specific brain regions. N2cG coated SAD-B19-ΔG has the ability to efficiently retrograde label the upstream network of specific brain regions along the axon terminal [18], while CVS-N2c-ΔG has lower cytotoxicity compared with SAD-B19-ΔG [10]. Therefore, if N2cG can endow the CVS-N2c-ΔG virus with efficient retrograde labeling, it will be more conducive to structural labeling and functional manipulation. To evaluate whether N2cG coated CVS-N2c-ΔG virus can achieve high-efficiency retrograde labeling projection neurons, 100 nl of the N2cG coated CVS-N2c-ΔG virus and CTB-488 (cholera toxin subunit B binding fluorescein 488, used to indicate the injection site) were mixed and injected into the ventral tegmental area (VTA) of C57BL/6J adult mice (Fig. 2A), and then local infection and the brain regions projecting to the VTA were imaged at 7 days post-injection (DPI). We found that N2cG coated CVS-N2c-ΔG virus only labeled a small number of neurons *in situ* (Fig. 2B), mainly retrogradely labeled the upstream brain area of VTA [19], including the somatomotor areas (MO), anterior cingulate area (ACA), medial preoptic area (MPOA), anterior hypothalamic nucleus (AHN), lateral habenula (LHb), lateral hypothalamic area (LHA), zona incerta (ZI), dorsal raphe nucleus (DR), and parabrachial nucleus (PB), among others (Fig. 2C), indicating the N2cG coated CVS-N2c-ΔG virus allows efficient retrograde access to projection neurons.

**Fig. 2.**
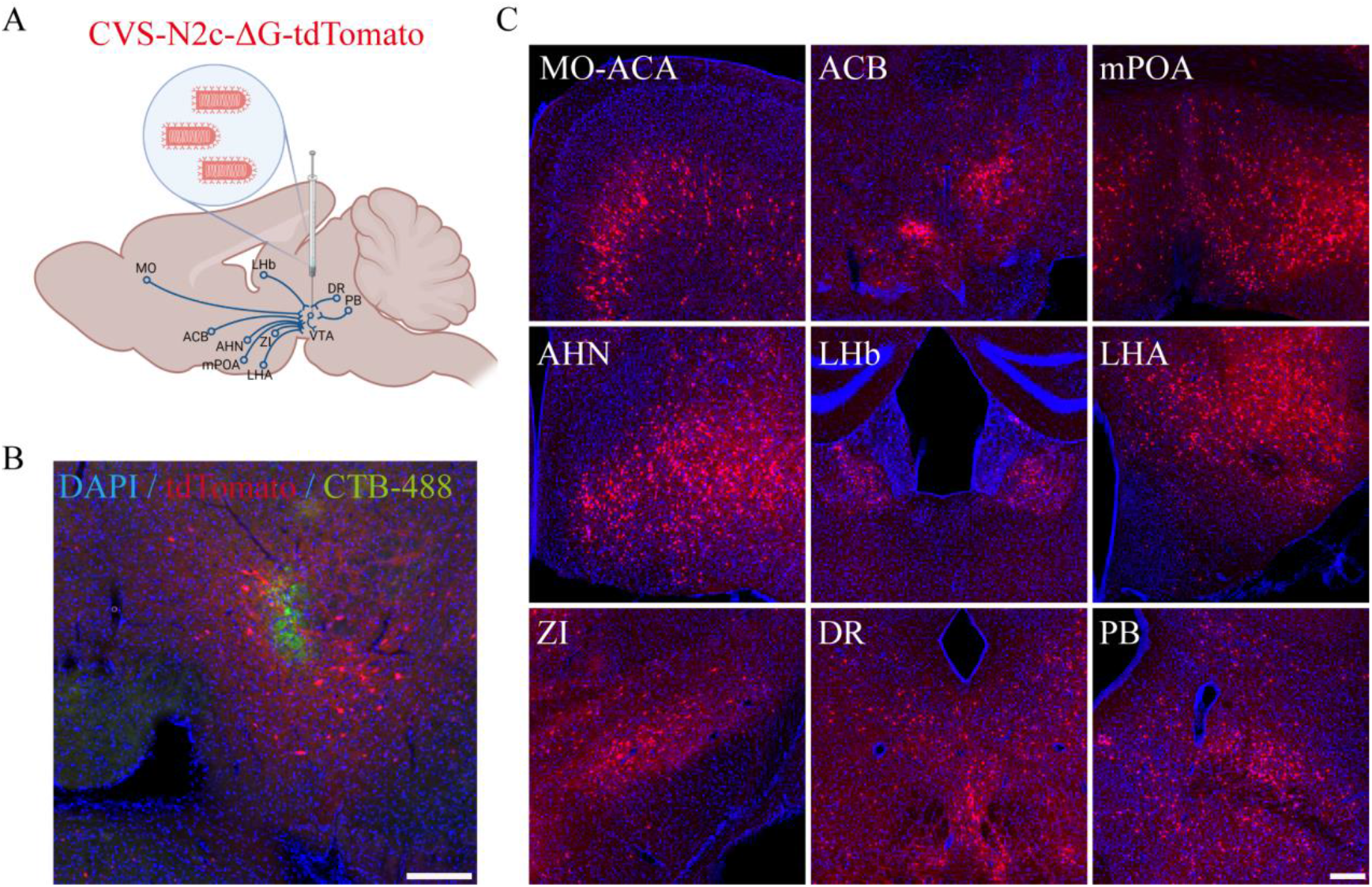
Efficient retrograde labeling with N2cG coated CVS-N2c-ΔG virus. (A) Schematic diagram of retrograde infection via N2cG coated CVS-N2c-ΔG virus. (B) N2cG coated CVS-N2c-ΔG virus could infect a small amount of neurons in injection site VTA. CTB-488 dye (green) was used to indicate the injection site. (C) N2cG coated CVS-N2c-ΔG virus could efficiently retrograde infect the upstream brain regions projecting to VTA, including the somatomotor areas (MO), anterior cingulate area (ACA), medial preoptic area (MPOA), anterior hypothalamic nucleus (AHN), lateral habenula (LHb), lateral hypothalamic area (LHA), zona incerta (ZI), dorsal raphe nucleus (DR), and parabrachial nucleus (PB), among others. Scale bars: 100 μm.

### Transduction efficiency of CVS-N2c-ΔG virus in VTA/SNc to DLS pathway

In order to exhibit that N2cG coated CVS-N2c-ΔG virus allows efficiently retrograde access to projection neurons difficult to label with other tools, we compared its efficiency with another retrograde viral tracers, rAAV9-Retro, which can retrogradely infect projection neurons with an efficiency comparable to that of AAV2-Retro [20]. AAV2-Retro and rAAV9-Retro exhibit robust retrograde functionality in certain neural circuits, but they have brain region selectivity, and have weak labeling efficiency in projection neurons from the ventral tegmental area and substantia nigra pars compacta (VTA/SNc) to dorsal lateral striatum (DLS) [20–22]. N2cG coated CVS-N2c-ΔG-tdTomato and rAAV9-Retro-CAG-EGFP were mixed (volume ratio of 1:1, 200 nL per mouse) and injected into the CPu (DLS) of C57BL/6J adult mice (Fig. 3A), and then local infection and the VTA/SNc region projecting to the DLS were imaged at 14 days post-injection (DPI). We found that rAAV9-Retro showed weak EGFP expression in VTA/SN. In contrast, N2cG coated CVS-N2c-ΔG robustly drove tdTomato expression in projection neurons in VTA/SNc (Fig. 3B), demonstrating its efficient retrograde transport in VTA/SNc to DLS pathway. Importantly, significant statistical diference between rAAV9-Retro and CVS-N2c-ΔG was found in the total number of positive neurons in VTA/SNc (Fig. 3C, 38.33 ± 0.88 for rAAV9-Retro, 262.30 ± 4.06 for CVS-N2c-ΔG; P < 0.0001). These results indicate that the N2cG coated CVS-N2c-ΔG allows efficient retrograde access to projection neurons unaddressed by rAAV9-Retro, and the efficiency is 6 folds higher than that of rAAV9-Retro.

**Fig. 3.**
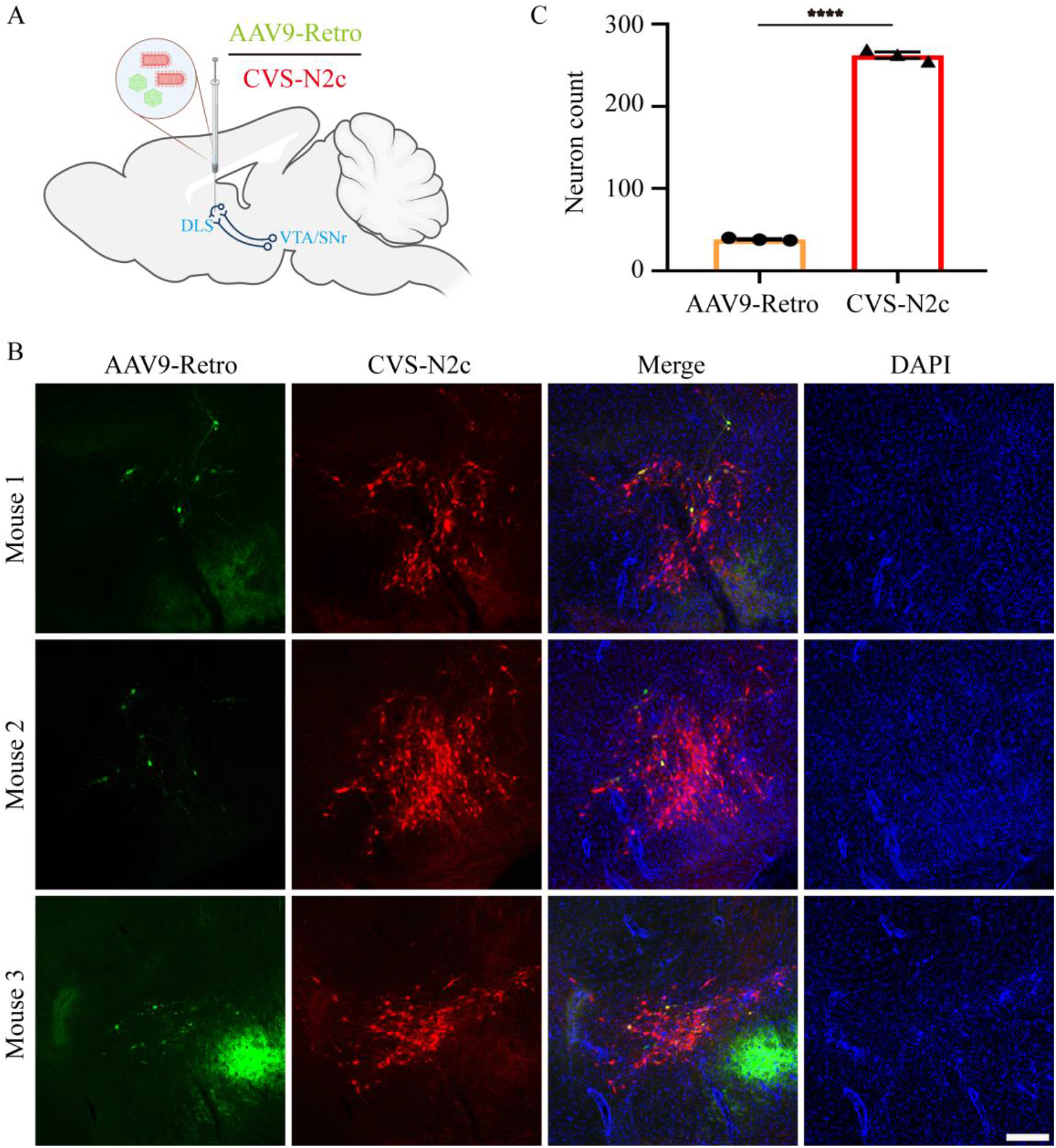
CVS-N2c-ΔG virus for efficient transduction in VTA/SNc to DLS pathway. (A) Schematic of CVS-N2c-ΔG-tdTomato and rAAV9-Retro-CAG-EGFP injections in VTA/SNc to DLS pathway. (B) Representative images of CVS-N2c-ΔG-tdTomato (red) and rAAV9-Retro-CAG-EGFP (green) in VTA/SNc. (C) Quantification of positive cells of viruses in VTA/SNc. Statistical values are indicated as mean ± SEM. Significant differences are expressed by the p value. *P<0.05, **P<0.01, ***P<0.001, ****P<0.0001, ns, no significant difference. Scale bars: 200 μm.

### Establishment and efficiency comparison of retrograde trans-mono-synaptic systems

Rabies virus strains from different sources combined with different glycoproteins may have different trans-synaptic efficiency or other infection tropism. The codon optimized chimeric glycoprotein (oG) derived from Pasteur strain of rabies virus can greatly improve trans-mono-synaptic tracing efficiency of SAD-B19-ΔG [11], so the retrograde trans-mono-synaptic system based on SAD-B19-ΔG/oG has been widely used to track the upstream input of neural networks. However, whether oG can enhance CVS-N2c-ΔG trans-mono-synaptic spread efficiency is still unknown. Therefore, it is necessary to compare different retrograde trans-mono-synaptic systems, as shown in Fig. 4, which mainly include rabies virus systems with deletion of glycoproteins and adeno-associated virus helper virus systems that compensate TVA for specific infection and glycoproteins for trans-mono-synaptic tracing. To verify whether oG can enhance CVS-N2c-ΔG trans-mono-synaptic spread efficiency, we used two helper viruses (AAVs) introduced in trans, one to complement the oG and the other to express TVA. They were mixed and injected into the ventral hippocampal region (vHPC) of Thy1-Cre transgenic mice. After 3 weeks of infection, CVS-N2c-ΔG and SAD-B19-ΔG was injected at the same site respectively. Seven days later, the brain slices were processed and imaged by slide scanner (Fig. 5A). We found that a number of nuclear GFP and dsRed neurons were co-labeled (starter cells) within vHPC (Fig. 5B), and rabies viruses could efficiently trans-mono-synaptic transduce the contralateral ventral hippocampal region (Fig. 5C and Fig. 5D). The trans-mono-synaptic tracing efficiency of rabies was evaluated through the convergence index, which is calculated as the number of dsRed^+^ input neurons divided by the number of GFP^+^ dsRed^+^ starter neurons [11]. Results showed that the trans-mono-synaptic efficiency of oG-mediated CVS-N2c-ΔG was 2-3 fold higher than that of oG-mediated SAD-B19-ΔG (Fig. 5E, CVS-N2c-ΔG/oG: 1.87 ± 0.16; SAD-B19-ΔG/oG: 0.60 ± 0.06; P = 0.0014).

**Fig. 4.**
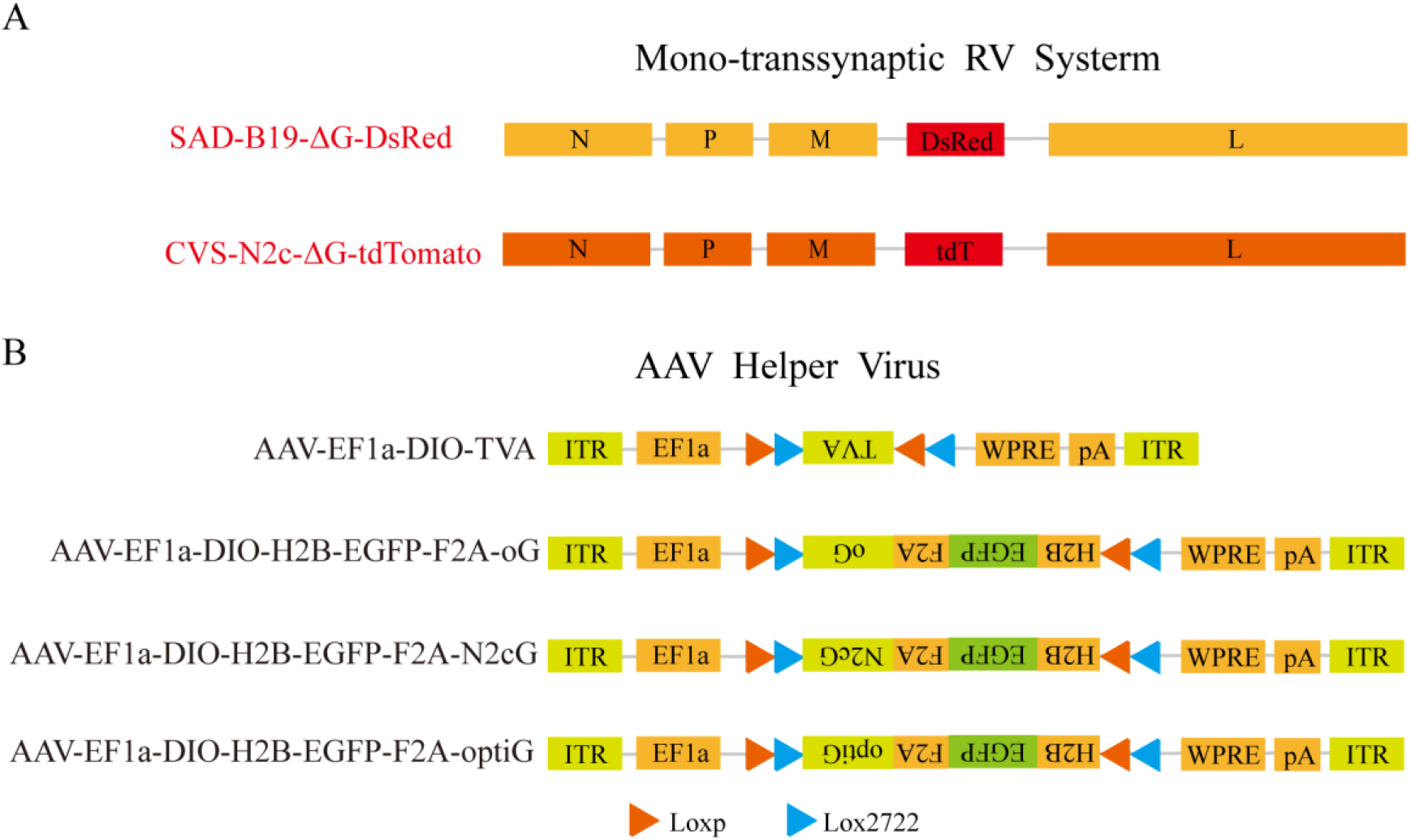
Establishment of retrograde trans-mono-synaptic systems. (A) Glycoprotein (RVG) deleted rabies virus systems: CVS-N2c-ΔG and SAD-B19-ΔG viral vectors for retrograde trans-mono-synaptic tracing. (B) Helper virus systems based on adeno-associated viruses that compensate TVA for cell-type specific infection of EnvARVG pseudotyped rabies viruses and glycoproteins for trans-mono-synaptic tracing. Different rabies virus systems and different helper virus systems can be combined into various retrograde trans-mono-synaptic systems. “tdTomato” is abbreviated as “tdT”‘ and optiG is codon-optimized N2cG. LoxP and Lox2722 are elements recognized and cleaved by Cre recombinase.

**Fig. 5.**
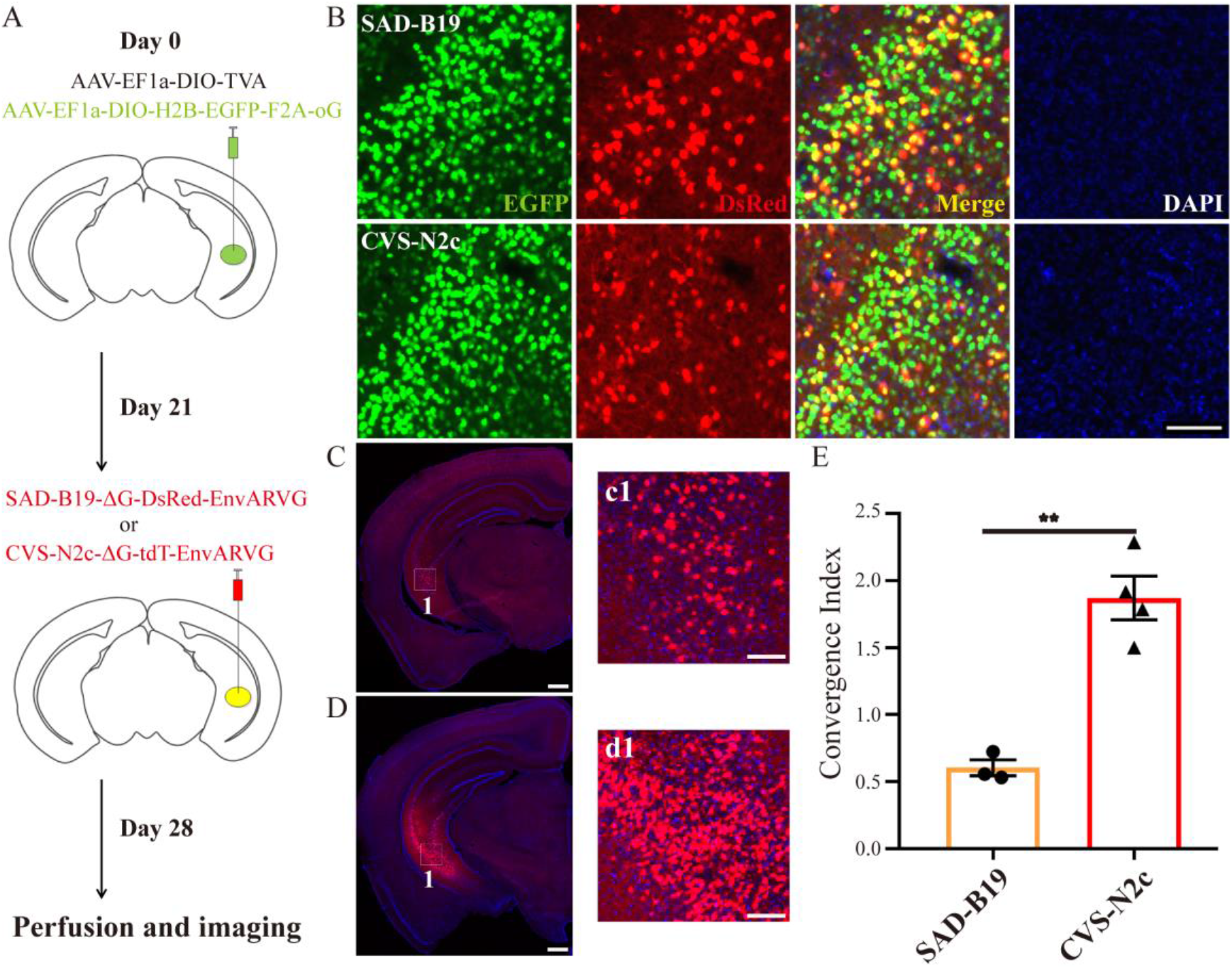
Comparison of retrograde trans-mono-synaptic efficiency following trans-complementation with oG. (A) Schematic diagram of virus injection for trans-mono-synaptic tracing. The adeno-associated viruses carrying Cre-dependent oG and TVA were injected into the ventral hippocampal region (vHPC) of Thy1-CRE transgenic mice. After 3 weeks, EnvARVG pseudotyped CVS-N2c-ΔG and SAD-B19-ΔG were injected at the same site respectively. After 1 week, brain slices were processed and imaged by slide scanner. (B) Starter cells at injection site. The green fluorescence signals of oG could be co-labeled with the red fluorescence signals of RV. (C) Monosynaptic input neurons in contralateral vHPC labeled by oG mediated SAD-B19-ΔG spread. c1 is a partial enlarged view of figure C. (D) Monosynaptic input neurons in contralateral vHPC labeled by oG mediated CVS-N2C-ΔG spread. d1 is a partial enlarged view of figure D. (E) Convergence indices for long-distance input in contralateral vHPC. Statistical values are indicated as mean ± SEM. Significant differences are expressed by the p value. *P<0.05, **P<0.01, ***P<0.001, ns, no significant difference. Scale bars: 100 μm for figure B/c1/d1; 500 μm for figure C/D.

Using the same method, we compared the trans-mono-synaptic efficiency of CVS-N2c-ΔG/oG and CVS-N2c-ΔG/N2cG. The schematic diagram of virus injection is shown in Fig. S2A, a number of starter cells were labeled in vHPC (Fig. S2B), and CVS-N2c-ΔG/N2cG could also efficiently trans-mono-synaptic transduce the contralateral ventral hippocampal region (Fig. S2C). Quantitative analysis showed that the retrograde trans-mono-synaptic efficiency of CVS-N2c-ΔG/oG was equivalent to that of CVS-N2c-ΔG/N2cG (Fig. S2D, CVS-N2c-ΔG/oG: 1.87 ± 0.16; CVS-N2c-ΔG/N2cG: 1.95 ± 0.15; P = 0.7337). However, oG is a codon optimized chimeric glycoprotein. Whether optimized N2cG (optiG) can further improve the trans-mono-synaptic efficiency of the CVS-N2c-ΔG virus is still unknown. We compared the trans-mono-synaptic efficiency of CVS-N2c-ΔG/N2cG and CVS-N2c-ΔG/optiG using D2R-Cre transgenic mice. The viruses were injected into the CPu region of D2R-Cre transgenic mice (Fig. S3A). A number of co-labeled signals could be observed at the injection site (Fig. S3B), and both CVS-N2c-ΔG/N2cG and CVS-N2c-ΔG/optiG could retrograde trans-mono-synaptic label a large number of neurons in the cortex (Fig. S3C and Fig. S3D), which was consistent with the previous report [10]. In addition, both trans-mono-synaptic systems can also efficiently retrograde target the amygdala and thalamus (Fig. S4A and Fig. S4C), the main inputs of CPu area. Through quantitative analysis, we found that the convergence indices had no significant difference in both amygdala (Fig. S4B, BLA, N2cG: 1.01 ± 0.37; optiG: 0.80 ± 0.19; P = 0.6211) and thalamus (Fig. S4D, TH, N2cG: 3.96 ± 0.81; optiG: 4.98 ± 1.56; P = 0.5850), respectively, indicating that optiG could not improve the trans-mono-synaptic efficiency of CVS-N2c-ΔG.

### CVS-N2c-ΔG virus for monitoring neural activity

While CVS-N2c-ΔG exhibits reduced toxicity to neurons compared with SAD-B19-ΔG vaccine strain, and can be used for long-term neuronal activity monitoring of labeled circuit in vivo, it still has some neurotoxicity, and the time for monitoring neural activity is 17 days longest tested [10]. Therefore, it is essential to evaluate the functional characteristics of viruses produced by a new method. To evaluate whether projection neurons transduced with the N2cG coated CVS-N2c-ΔG vector retained the properties for monitoring neural activity, and to determine the time window it maintains for function detection, we established CVS-N2c-ΔG-GCaMP6s viral vector and conducted in vivo response monitoring of calcium transients of reward circuits at different time points. The projection neurons from the ventral tegmental area (VTA) to nucleus accumbens (NAc) are involved in “reward circuits” [23], we performed fiber photometry in this projection pathway (Fig. 6A), and used 5% sugar water as reward. We injected N2cG coated CVS-N2c-ΔG-GCaMP6s vector into NAc and detected the change of calcium signals in VTA when mice were rewarded with sugar water (Fig. 6A, B). We found that the projection pathway could be labeled by GCaMP6s (Fig. 6C, D) and activated only when mice licked sugar water at different time points (Fig. 6E-G, 7, 14 and 21 days post infection). These results indicate that the N2cG coated CVS-N2c-ΔG can be used for monitoring neural activity of projection neurons, and the time window can be maintained for at least 21 days.

**Fig. 6.**
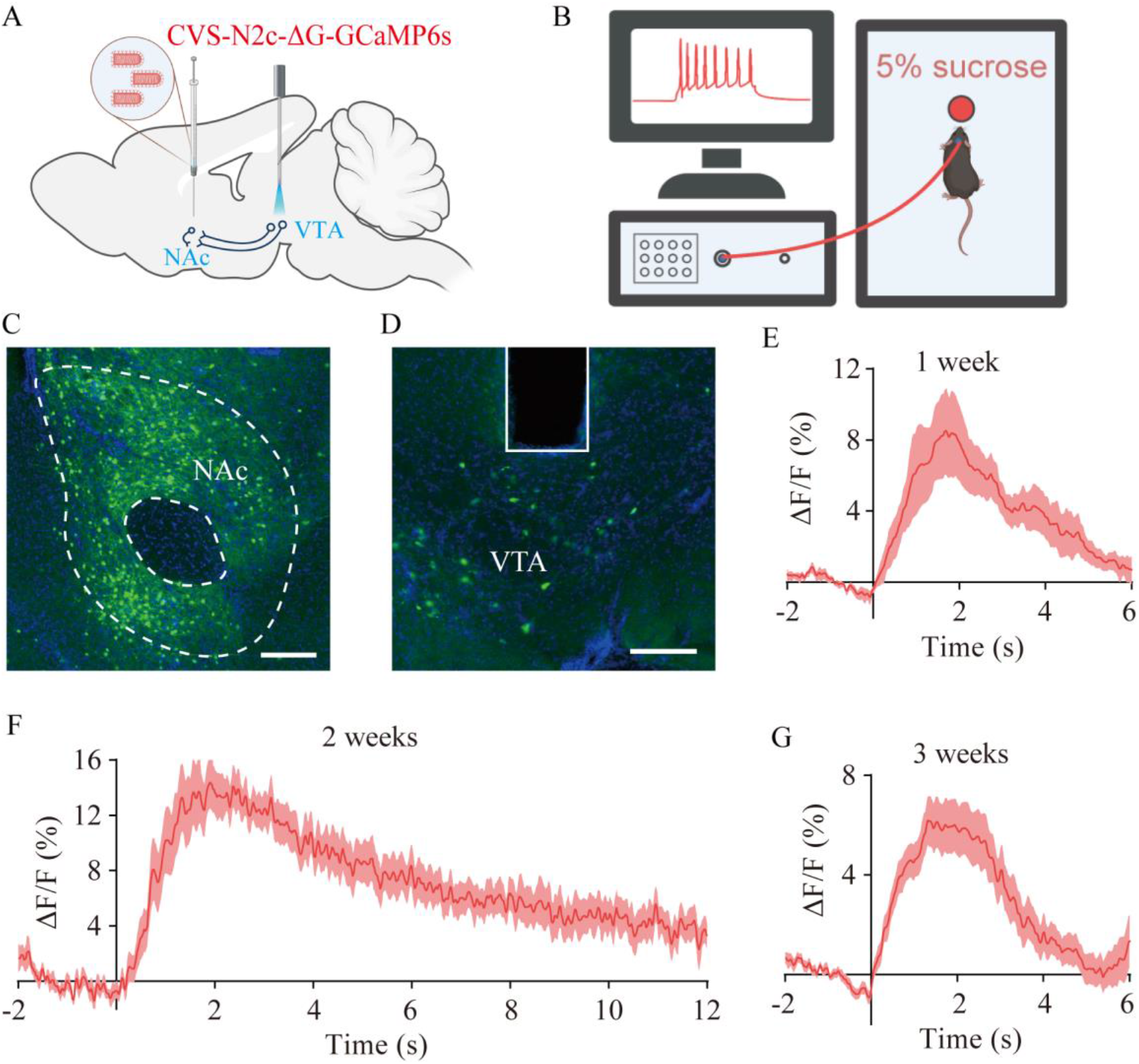
Neuronal activity monitoring in vivo with CVS-N2c-ΔG virus. (A) Schematic diagram of virus injection and optical fiber implantation. N2cG coated CVS-N2c-ΔG-GCaMP6s vector was injected into NAc and optical fiber was implanted into VTA. (B) Schematic diagram of reward behavior experiment and calcium transient monitoring. (C) Fluorescence signals of neurons in NAc labeled by CVS-N2c-ΔG-GCaMP6s. (D) Fluorescence signals of projection neurons in VTA labeled by CVS-N2c-ΔG-GCaMP6s. (E-G) Average of ΔF/F signals from reward behavior trials. Scale bars: 200 μm.

### CVS-N2c-ΔG expressing recombinase for transgene recombination

Recombinant enzyme mediated gene expression or manipulation plays an important role in the study of neural circuit structure and function [16]. In order to verify whether the CVS-N2c-ΔG virus produced by using the new method can effectively express recombinant enzyme for the recombination of Cre or Flpo induced genes. For Cre-mediated recombination, Cre-conditional rAAV expressing EGFP (rAAV-DIO-EGFP) was injected into the primary motor cortex (M1) of C57BL/6J adult mice, followed by injection of CVS-N2c-ΔG-mCherry-2A-Cre at the contralateral M1 (Fig. 7A); For Flpo-mediated recombination, Flpo-conditional rAAV expressing EGFP (rAAV-FDIO-EGFP) was injected into the primary motor cortex (M1) of C57BL/6J adult mice, followed by injection of CVS-N2c-ΔG-mCherry-2A-Flpo at the contralateral M1 (Fig. 7C). We found that both CVS-N2c-ΔG-mCherry-2A-Cre and CVS-N2c-ΔG-mCherry-2A-Flpo could transport retrogradely and drive EGFP expression of AAV, and the green fluorescence signals co-labeled with the red fluorescence signals (Fig. 7B and Fig. 7D). These results indicate that CVS-N2c-ΔG virus produced by using the new method can express sufficient recombinases for efficient transgene recombination, consistent with previous reports.

**Fig. 7.**
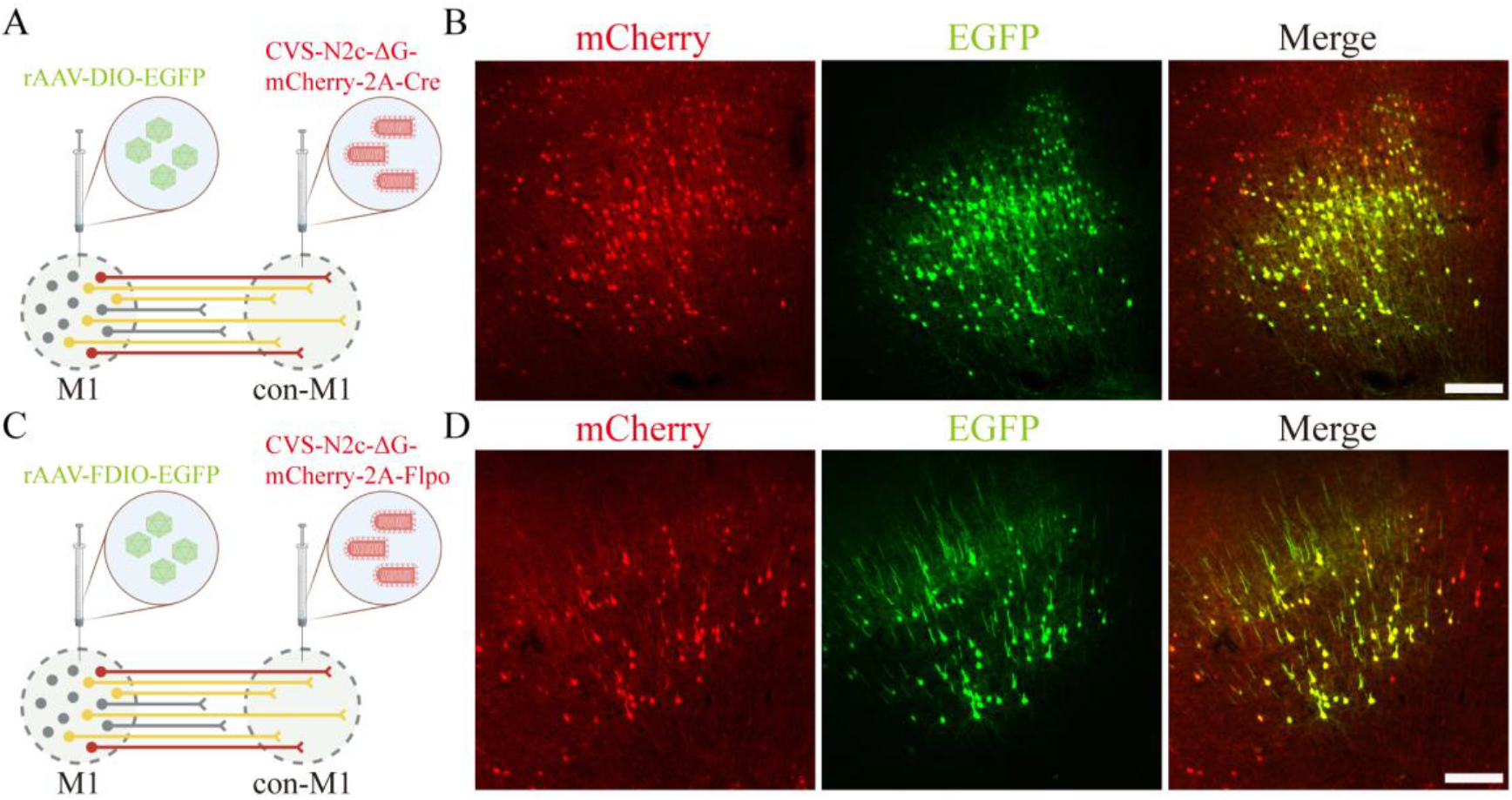
Cre- or Flp-dependent recombination with CVS-N2c-ΔG virus. (A) Schematic of virus injection and recombination by CVS-N2c-ΔG-mCherry-2A-Cre. Cre-conditional rAAV expressing EGFP (rAAV-DIO-EGFP) was injected into the primary motor cortex (M1) of C57BL/6J adult mice, followed by injection of CVS-N2c-ΔG-mCherry-2A-Cre at the contralateral M1 (con-M1). (B) Images of M1 indicating Cre-mediated recombination and expression in retrogradely infected neurons. (C) Schematic of virus injection and recombination by CVS-N2c-ΔG-mCherry-2A-Flpo. Flpo-conditional rAAV expressing EGFP (rAAV-FDIO-EGFP) was injected into the primary motor cortex (M1) of C57BL/6J adult mice, followed by injection of CVS-N2c-ΔG-mCherry-2A-Flpo at the contralateral M1. (D) Images of M1 indicating Flpo-mediated recombination and expression in retrogradely infected neurons. Scale bars: 200 μm.

## Discussion

Neurotropic virus tracers, especially those with low toxicity or high efficient tracing, contribute to the analysis of the anatomical structure and function of neural circuits [10, 16]. The retrograde trans-mono-synaptic system based on the transformation of rabies virus CVS-N2c strain has reduced cytotoxicity and improved trans-synaptic efficiency compared with the traditional SAD-B19 trans-mono-synaptic system [10], but it has not been further popularized in the field of neuroscience, mainly due to the defect of difficult preparation and low titer. To overcome these shortcomings, a new production system was established for rapid rescue and preparation of CVS-N2c-ΔG virus. The CVS-N2c-ΔG virus allows efficient retrograde access to projection neurons unaddressed by rAAV9-Retro, and maintains excellent performances for retrograde trans-mono-synaptic targeting, functional monitoring and transgene recombination.

CVS-N2c is highly neuroinvasive and can be rapidly transduced in the nervous system [24]. The previously reported preparation process of RVG deleted CVS-N2c virus mainly uses the cell line established by Neuro2a, which is not easy to culture [10]. It takes a long time to screen the cell line, and the amplification efficiency using its own glycoprotein is low, which often leads to low virus production efficiency. Alternative methods need to be found to produce RVG deleted CVS-N2c virus. However, high titer SAD-B19 virus is mainly produced by BHK-21 cells that are easy to culture and transduce [25]. Therefore, the cell line derived from BHK-21 may be able to quickly rescue and expand CVS-N2c-ΔG. In order to prove this phenomenon, CVS-N2c-ΔG was rescued in B7GG cell line, and obvious virus fluorescence signals could be observed on the fifth day, and it could be amplified rapidly. The supernatant titer can reach the level of SAD-B19-ΔG, and the preparation time is 10 ~ 14 days, indicating that B7GG cell line can quickly rescue and prepare CVS-N2c-ΔG virus. When infected with BHK-N2cG cell line, CVS-N2c-ΔG virus could also be amplified, but the titer of supernatant was significantly lower than that produced by B7GG cell line, indicating that B7GG cell line is more suitable for high titer production of CVS-N2c-ΔG virus. In addition, the supernatant titers of EnvARVG pseudotyped CVS-N2c-ΔG and SAD-B19-ΔG viruses prepared by BHK-EnvARVG cell line had no significant difference, indicating that BHK-EnvARVG cell line can prepare EnvARVG pseudotyped CVS-N2c-ΔG virus with high titer, which is inconsistent with the previously reported low production efficiency of CVS-N2c-ΔG pseudotyped with EnvARVG [10], which may be related to Neuro2a cell line that are not easy to culture. The production time of EnvARVG pseudotype CVS virus is 5 ~ 7 days. Therefore, this new method greatly shortens the preparation cycle of CVS-N2c-ΔG virus.

Some viral vectors have been used as retrograde tracers to label and manipulate neurons projected to a specific brain region. Sun et al. [26] and Zhu et al. [18] reported in detail the retrograde labeling efficiency of various viral tracers, including PRV, RV-B2C, AAV2-Retro and N2cG pseudotyped SAD-B19, and found that they have different retrograde labeling efficiency and brain region selectivity. The brain area targeting of N2cG pseudotyped SAD-B19 is more broad-spectrum than that of AAV2 retro, and the retrograde labeling efficiency of N2cG pseudotyped SAD-B19 is more than one order of magnitude higher than that of RV packaged with B19G, providing a valuable reference for the selection and use of tool viruses [18]. CVS-N2c has lower cytotoxicity than SAD-B19, therefore, if N2cG encapsulated CVS-N2c-ΔG virus permits efficient retrograde access to projection neurons, which will be more conducive to structural labeling and functional manipulation. We found that N2cG coated CVS-N2c-ΔG allows efficient retrograde access to projection neurons, and further expand its application in VTA/SNc to DLS pathway that unaddressed by rAAV9-Retro, and the efficiency is 6 folds higher than that of rAAV9-Retro. In future studies, we will use this viral tracer to trace and manipulate more neural circuits that cannot be resolved by other viral tracers.

The reported efficient retrograde trans-mono-synaptic systems mainly include CVS-N2c-ΔG/N2cG [10] and SAD-B19-ΔG/oG [11], but the trans-synaptic efficiency of CVS-N2c-ΔG mediated by oG and modified N2cG (optiG) is unknown. In order to answer this question, A variety of retrograde trans-mono-synaptic systems were established by combining different rabies virus systems with different sources of RVG, and the efficiency was compared in the nervous system of mice. Results showed that the trans-mono-synaptic efficiency of oG-mediated CVS-N2c-ΔG was 2-3 folds higher than that of oG-mediated SAD-B19-ΔG, but there was no difference between oG-mediated and N2cG-mediated CVS-N2c-ΔG system. In addition, codon modified N2cG (optiG) did not increase the efficiency of CVS-N2c-ΔG tracing. Therefore, retrograde trans-mono-synaptic system using oG-mediated or N2cG-mediated CVS-N2c-ΔG is conducive to analyze more comprehensive input network. It should be noted that other RVG from different sources not included here may also further improve the trans-mono-synaptic efficiency of the virus, more comparative studies are needed later. In addition, the trans-mono-synaptic efficiency of CVS-N2c-ΔG/oG is higher than that of SAD-B19-ΔG/oG, which may be because oG is easier to package CVS-N2c-ΔG virus, or CVS-N2c-ΔG virus is less toxic, which is more conducive to the maintenance of nerve cells and the trans-mono-synaptic transmission of CVS-N2c-ΔG virus. The relevant mechanism needs to be further studied. Then, oG can be constructed into lentivirus vector, packaged into lentivirus, infected with BHK-21 and other cells to establish a stable cell line, which may be useful to more efficiently package or prepare CVS-N2c-ΔG virus, and what are the labeling characteristics of the packaged pseudovirus in vivo? It is also a problem worthy of study.

Finally, we found that the CVS-N2c-ΔG produced by the new method can be used for monitoring neural activity of projection neurons, and the time window can be maintained for 3 weeks, which is at least consistent with the previous report (up to 17 days) [10]. In addition, CVS-N2c-ΔG virus produced using the new method can express sufficient recombinases for efficient transgene recombination, which is also consistent with previous reports [10]. Therefore, the virus produced by the new method does not affect its own function, paving the way for its further optimization and application in neural circuit.

In summary, this work provides improved production method and expanded application of CVS-N2c-ΔG virus, which will contribute to the popularization and application of CVS-N2c-ΔG virus for structural and functional studies of neural circuits.

## Materials and methods

### Establishment of stable cell lines for virus packaging

BHK-21 cells stably expressing either N2cG or EnvARVG (a chimeric protein made from EnvA and the tail of B19G) fused with nuclear localized EGFP were created. H2B-GFP-P2A-N2cG and H2B-GFP-P2A-EnvARVG fragments were cloned into lentivirus expression vector FUGW (addgene#14883) by homologous recombination kit (Vazyme company), and FUGW-H2B-GFP-P2A-N2cG and FUGW-H2B-GFP-P2A-EnvARVG vectors were obtained and transfected into HEK-293T with pMDLg/pRRE (addgene#12251), pRSV-Rev (addgene#12253), and pMD2.G (addgene#12259). Viral supernatants were collected at 48 and 72 hours post transfection and after filtering used to transduce BHK-21 cells at a multiplicity of infection (MOI) of 5. Three days post transduction, the green fluorescence ratio reached more than 90%, then the cells were passaged for 5 times and stored in liquid nitrogen. The resulting cell lines based on BHK-21 are named BHK-N2cG and BHK-EnvARVG.

### Vectors construction

The optiG was codon optimized from N2cG for M. musculus (Genscript). It was cloned into pAAV-Ef1a-DIO-H2B-GFP-P2A-N2cG plasmid (addgene#73476) to replace N2cG to obtain pAAV-Ef1a-DIO-H2B-GFP-P2A-optiG plasmid. To construct CVS-N2c-ΔG-GCaMP6s plasmid, calcium-sensitive probe GCaMP6s was synthesized and inserted into CVS-N2c-ΔG-tdTomato (addgene#73462) digested by the restriction enzymes SmaI and NheI (New England Biolabs).

### Rescue and preparation of rabies viral vectors

SAD-B19-ΔG-DsRed was rescued and prepared according to a previously reported method [17, 25]. B7GG cells were used for the rapid rescue and amplification of CVS-N2c-ΔG viruses. B7GG cells were cultured in good growth state, digested with 0.25% trypsin for 2 minutes, passaged to 6-well plates followed by adding 2 ML Dulbecco’s minimum essential media (DMEM) containing 10% fetal bovine serum (FBS), then cultured in 37 °C 5% CO2 incubator overnight. When the cell density in the 6-well plate reaches 80%, rescue was performed in the cells by co-transfection (Fugene 6 transfection reagent) of CVS-N2c-ΔG genomic plasmid, pCAG-B19P, pCAG-B19N, pCAG-B19L and pCAG-B19G. 3-5 days post transfection, fluorescence signals of the virus were observed, indicating that the virus was rescued successfully, then the cells were passaged to amplify CVS-N2c-ΔG virus pseudotyped with B19G in culture conditions of 35 °C and 3% CO2. The period of B19G pseudotyped CVS-N2c-ΔG virus rescue and amplification was 10 ~ 14 days. Amplification of N2cG coated CVS-N2c-ΔG virus was performed by adding amplified supernatant of B19G pseudotyped CVS-N2c-ΔG to BHK-N2cG cells. The production cycle of N2cG coated CVS-N2c-ΔG virus is 5 ~ 7 days. Amplification of EnvARVG pseudotyped CVS-N2c-ΔG virus was performed by adding amplified supernatant of B19G pseudotyped CVS-N2c-ΔG to BHK-EnvARVG cells. The production cycle of EnvARVG pseudotyped CVS-N2c-ΔG virus is 5 ~ 7 days.

The supernatants were collected and filtered with 0.22 μm membrane, then centrifuged at 50000×g for 2.5 h at 4 °C. The precipitation was suspended with 1 mL PBS and then concentrated and purified with 20% sucrose for the second time. The precipitation was suspended with appropriate amount of PBS. The titer was determined by 10 fold gradient dilution method (10^0^ ~ 10^-6^), and the virus was stored at – 80 °C until use.

Titers of N2cG-enveloped or B19G-pseudotyped viruses were tested using HEK-293T cells; Titers of EnvARVG-pseudotyped viruses were tested using HEK293T-TVA800 cells [27]. Titers of CVS-N2c-ΔG viruses (infectious units per mL, IU/mL) were 10^7^ – 10^8^ for N2cG-enveloped and 10^8^ – 10^9^ for B19G- and EnvARVG-pseudotyped. The rabies virus genome plasmids include CVS-N2c-ΔG-tdTomato (addgene#73462), CVS-N2c-ΔG-mCherry-2A-Cre (addgene#73472) and CVS-N2c-ΔG-mCherry-2A-Flpo (addgene#73471). These rabies viral vectors can be purchased from the BrainCase (ShenZhen, China).

### Production of adeno-associated viruses

AAV vectors were produced in HEK-293T cells cotransfected with pRep2Cap9 and pAdDeltaF6 (addgene#112867) using polyethylenimine (PEI-MAX), and then purified by iodixanol gradient ultracentrifugation [28]. The purified AAV vectors were titered by qPCR using the iQ SYBR Green Supermix kit (Bio–Rad). AAV vectors were stored at - 80 °C, and the titers were diluted to 2 x 10^12^ viral genomes/mL (VG/mL) with phosphate-buffered saline (PBS) before use, respectively. AAV vectors include rAAV-Ef1a-DIO-TVA (BrainCase), rAAV-Ef1a-DIO-H2B-GFP-P2A-oG (addgene#74289), rAAV-Ef1a-DIO-H2B-GFP-P2A-N2cG (addgene#73476) and rAAV-Ef1a-DIO-H2B-GFP-P2A-optiG. rAAV9-Retro-CAG-EGFP viral vector (1.0 x 10^13^ VG/mL) was purchased from the BrainCase (ShenZhen, China).

### Animals

Adult male (8–10 weeks old) C57BL/6J mice (Hunan SJA Laboratory Animal Company), Thy1-Cre and D2R-Cre transgenic mice were used for all experiments. Among them, Thy1-Cre transgenic mice were presented by professor Duan Shumin Laboratory (Zhejiang University); D2R-Cre transgenic mice were presented by researcher Xiong Zhiqi Laboratory (Institute of Neuroscience, Chinese Academy of Sciences). The mice were housed in the appropriate environment with a 12/12-h light/dark cycle, and water and food were supplied *ad libitum*. Through cross breeding with C57BL/6J mice in SPF level animal room, the offspring of transgenic mice identified as positive by gene identification were used for the experiments. All surgical and experimental procedures were performed in accordance with the guidelines formulated by the Animal Care and Use Committee of the Innovation Academy for Precision Measurement Science and Technology, Chinese Academy of Sciences.

### Virus injection

All the experiments related to AAV and RV viruses were performed in Biosafety Level-2 (BSL-2) laboratory. The stereotactic injection coordinates were selected according to Paxinos and Franklin’s *The Mouse Brain in Stereotaxic Coordinates*, 4th edition [29]. The stereotactic coordinates for VTA were as follows: anterior-posterior-axis (AP): - 3.20 mm; medial-lateral-axis (ML): ± 0.45 mm; dorsal-ventral-axis (DV): - 4.30 mm from bregma. The stereotactic coordinates for vHPC were as follows: AP: - 3.16 mm; ML: ± 2.95 mm; and DV: - 4.10 mm from bregma. The stereotactic coordinates for CPu were as follows: AP: + 0.38 mm; ML: ± 2.00 mm; DV: - 3.50 mm from bregma. Eight-to ten-week-old C57BL/6J mice and transgenic mice (20–25 g) were used for virus injection, and the standard injection process was performed as previously reported [18]. In the test of transsynaptic tracing, EnvARVG-pseudotyped rabies virus (3 x 10^8^ IU/mL, 100 nL per mouse) was injected at the same site 3 weeks after the injection of AAV helper viruses (TVA: RVG, volume ratio of 1:2, 150 nL per mouse), and the mice were sacrificed at 7 days post-injection using the conventional cardiac perfusion method. For retrograde tracing, the used titer of N2cG coated CVS-N2c-ΔG-tdTomato was 1 x 10^8^ IU/mL, and the used titer of N2cG coated CVS-N2c-ΔG-GCaMP6s was 3 x 10^7^ IU/mL.

### Slice preparation and imaging

Slice preparation and imaging were accomplished according to previously reported methods [20]. After soaked with 4% paraformaldehyde solution overnight, dehydration of mice brains was performed with 30% sucrose solution at 37 °C, then the coronal sections (thickness of 40 μm) of brains were completed by using a microtome (Thermo Fisher Scientific), and collected in anti-freeze fluid at 200-μm intervals. The brain slices were washed 3 times with phosphate-buffered saline (PBS) for 5 minutes each time. After 4’,6-diamidino-2-phenylindole (DAPI) staining (diluted at 1:3000) for 10 minutes, the brain slices were washed 3 times with PBS for 5 minutes each time, and applied neatly on microscope slides, followed by sealing with 70% glycerol. Imaging was performed using an Olympus VS120 Slide Scanner microscope (Olympus, Japan).

### Fiber photometry for neural activity recording

N2cG coated CVS-N2c-ΔG-GCaMP6s (150 nL) was injected into the NAc area, and optical fiber (core diameter: 200 μm, numerical aperture: 0.37, Inper, China) was implanted into the VTA area, then the change of calcium signals was detected in VTA when mice were rewarded with sugar water at different time points (7, 14 and 21 days post infection). Before recording the change of calcium signals, the mice were touched gently for 5 min/day at least 3 days, then habituated to a chamber and the fiber patch cord (20×20×22 cm) for 10 min. All mice were water-deprived for more than 24 h until they were placed in the chamber equipped with a cup, which was filled with 5 % (w/v) sucrose solution. Then the mice were tested when given sucrose rewards. Calcium transients in VTA during reward behavior were recorded by using a fiber photometry system (ThinkerTech, Nanjing, China) to excite GCaMP6s at 470 nm wavelength. For each trial, fluorescence signals from GCaMP6s were normalized by calculating z-scores as ΔF/F signals, where the mean and standard error of mean (SEM) was taken from a 2-s baseline acquisition period preceding sugar water delivery. Data were analyzed using MATLAB (MathWorks) code and Prism (GraphPad software).

## Abbreviations

AAV: adeno-associated virus
PBS: phosphate-buffered saline
VG: viral genomes
IU: infectious units
DAPI: 4’,6-diamidino-2-phenylindole
CPu: caudate putamen (striatum)
VTA: ventral tegmental area
vHPC: ventral hippocampus
MO: somatomotor areas
ACA: anterior cingulate area
MPOA: medial preoptic area
AHN: anterior hypothalamic nucleus
LHb: lateral habenula
LHA: lateral hypothalamic area
ZI: zona incerta
DR: dorsal raphe nucleus
PB: parabrachial nucleus
SNr: substantia nigra pars compacta
BLA: basolateral amygdalar nucleus
TH: thalamus.

## Authors’ contributions

KL and FX contributed to the study idea and design; KL and FX contributed to funding acquisition and resources; KL, LL, WM, XY and NL performed the experiments and data acquisition; KL, LL, WM and ZH performed the data analysis; KL and FX drafted the manuscript and contributed to its review and editing. All authors read and approved the final manuscript.

## Acknowledgements

This work was supported by the National Natural Science Foundation of China (32100899, 31830035, 31771156, 21921004), the Key-Area Research and Development Program of Guangdong Province (2018B030331001), the Strategic Priority Research Program of the Chinese Academy of Sciences (XDB32030200) and the Shenzhen Key Laboratory of Viral Vectors for Biomedicine (ZDSYS20200811142401005). Schematic diagrams of the process in the figures (figure 1, 2, 3, 6, 7) of this article were created with BioRender.com.

## References

1. Insel TR. Rethinking schizophrenia. Nature. 2010;468(7321):187–193.

2. Li J, Liu T, Dong Y, Kondoh K and Lu Z. Trans-synaptic Neural Circuit-Tracing with Neurotropic Viruses. Neurosci Bull. 2019;35(5):909–920.

3. Smith BN, Banfield BW, Smeraski CA, Wilcox CL, Dudek FE, Enquist LW, et al. Pseudorabies virus expressing enhanced green fluorescent protein: A tool for in vitro electrophysiological analysis of transsynaptically labeled neurons in identified central nervous system circuits. Proceedings of the National Academy of Sciences of the United States of America. 2000;97(16):9264–9269.

4. Saleeba C, Dempsey B, Le S, Goodchild A and McMullan S. A Student’s Guide to Neural Circuit Tracing. Front Neurosci. 2019;13:897.

5. Lin K, Zhong X, Ying M, Li L, Tao S, Zhu X, et al. A mutant vesicular stomatitis virus with reduced cytotoxicity and enhanced anterograde trans-synaptic efficiency. Mol Brain. 2020;13(1):45.

6. Wickersham IR, Lyon DC, Barnard RJ, Mori T, Finke S, Conzelmann KK, et al. Monosynaptic restriction of transsynaptic tracing from single, genetically targeted neurons. Neuron. 2007;53(5):639–47.

7. Wickersham IR, Finke S, Conzelmann KK and Callaway EM. Retrograde neuronal tracing with a deletion-mutant rabies virus. Nat Methods. 2007;4(1):47–9.

8. Xiao Q, Zhou X, Wei P, Xie L, Han Y, Wang J, et al. A new GABAergic somatostatin projection from the BNST onto accumbal parvalbumin neurons controls anxiety. Mol Psychiatry. 2021;26(9):4719–4741.

9. Schwarz LA, Miyamichi K, Gao XJ, Beier KT, Weissbourd B, DeLoach KE, et al. Viral-genetic tracing of the input-output organization of a central noradrenaline circuit. Nature. 2015;524(7563):88–92.

10. Reardon TR, Murray AJ, Turi GF, Wirblich C, Croce KR, Schnell MJ, et al. Rabies Virus CVS-N2c(DeltaG) Strain Enhances Retrograde Synaptic Transfer and Neuronal Viability. Neuron. 2016;89(4):711–24.

11. Kim EJ, Jacobs MW, Ito-Cole T and Callaway EM. Improved Monosynaptic Neural Circuit Tracing Using Engineered Rabies Virus Glycoproteins. Cell Rep. 2016;15(4):692–699.

12. Szőnyi A, Sos KE, Nyilas R, Schlingloff D, Domonkos A, Takács VT, et al. Brainstem nucleus incertus controls contextual memory formation. Science. 2019;364(6442).

13. Szőnyi A, Zichó K, Barth AM, Gönczi RT, Schlingloff D, Török B, et al. Median raphe controls acquisition of negative experience in the mouse. Science. 2019;366(6469).

14. Hafner G, Witte M, Guy J, Subhashini N, Fenno LE, Ramakrishnan C, et al. Mapping Brain-Wide Afferent Inputs of Parvalbumin-Expressing GABAergic Neurons in Barrel Cortex Reveals Local and Long-Range Circuit Motifs. Cell Rep. 2019;28(13):3450–3461.e8.

15. Ciabatti E, Gonzalez-Rueda A, Mariotti L, Morgese F and Tripodi M. Life-Long Genetic and Functional Access to Neural Circuits Using Self-Inactivating Rabies Virus. Cell. 2017;170(2):382–392 e14.

16. Chatterjee S, Sullivan HA, MacLennan BJ, Xu R, Hou Y, Lavin TK, et al. Nontoxic, double-deletion-mutant rabies viral vectors for retrograde targeting of projection neurons. Nat Neurosci. 2018;21(4):638–646.

17. Osakada F, Mori T, Cetin AH, Marshel JH, Virgen B and Callaway EM. New rabies virus variants for monitoring and manipulating activity and gene expression in defined neural circuits. Neuron. 2011;71(4):617–31.

18. Zhu X, Lin K, Liu Q, Yue X, Mi H, Huang X, et al. Rabies Virus Pseudotyped with CVS-N2C Glycoprotein as a Powerful Tool for Retrograde Neuronal Network Tracing. Neurosci Bull. 2020;36(3):202–216.

19. Montardy Q, Zhou Z, Lei Z, Liu X, Zeng P, Chen C, et al. Characterization of glutamatergic VTA neural population responses to aversive and rewarding conditioning in freely-moving mice. Science Bulletin. 2019;64(16):1167–1178.

20. Lin K, Zhong X, Li L, Ying M, Yang T, Zhang Z, et al. AAV9-Retro mediates efficient transduction with axon terminal absorption and blood-brain barrier transportation. Mol Brain. 2020;13(1):138.

21. Tervo DG, Hwang BY, Viswanathan S, Gaj T, Lavzin M, Ritola KD, et al. A Designer AAV Variant Permits Efficient Retrograde Access to Projection Neurons. Neuron. 2016;92(2):372–382.

22. Li SJ, Vaughan A, Sturgill JF and Kepecs A. A Viral Receptor Complementation Strategy to Overcome CAV-2 Tropism for Efficient Retrograde Targeting of Neurons. Neuron. 2018;98(5):905–917 e5.

23. Mohebi A, Pettibone JR, Hamid AA, Wong JT, Vinson LT, Patriarchi T, et al. Dissociable dopamine dynamics for learning and motivation. Nature. 2019;570(7759):65–70.

24. Bostan AC, Dum RP and Strick PL. The basal ganglia communicate with the cerebellum. Proc Natl Acad Sci U S A. 2010;107(18):8452–6.

25. Osakada F and Callaway EM. Design and generation of recombinant rabies virus vectors. Nat Protoc. 2013;8(8):1583–601.

26. Sun L, Tang Y, Yan K, Yu J, Zou Y, Xu W, et al. Differences in neurotropism and neurotoxicity among retrograde viral tracers. Mol Neurodegener. 2019;14(1):8.

27. Narayan S, Barnard RJ and Young JA. Two retroviral entry pathways distinguished by lipid raft association of the viral receptor and differences in viral infectivity. J Virol. 2003;77(3):1977–83.

28. Chen YH, Keiser MS and Davidson BL. Adeno-Associated Virus Production, Purification, and Titering. Curr Protoc Mouse Biol. 2018;8(4):e56.

29. Paxinos G and Franklin K. (2012) Paxinos and Franklin’s the Mouse Brain in Stereotaxic Coordinates, Amsterdam: Elsevier/Academic Press.

