## Supplemental figures for "Improved production and expanded application of CVS-N2c-ΔG virus for retrograde tracing"

### Supplemental data

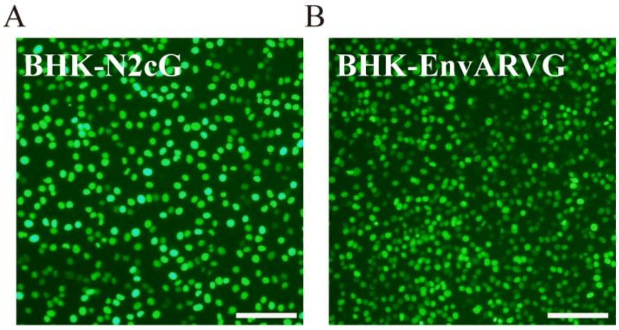


**Fig. S1 Establishment of stable cell lines.** (A) "BHK-N2cG" cell line, based on BHK-21 cells stably expressing the CVS-N2c glycoprotein (N2cG), along with the nuclear-localized EGFP, used for the amplification of N2cG coated CVS-N2c-ΔG viruses; (B) "BHK-EnvARVG" cell line, based on BHK-21 cells stably expressing the EnvA and B19G chimeric glycoprotein (EnvARVG), along with the nuclear-localized EGFP, used for the amplification of EnvARVG coated CVS-N2c-ΔG viruses. Scale bars: 100 μm.

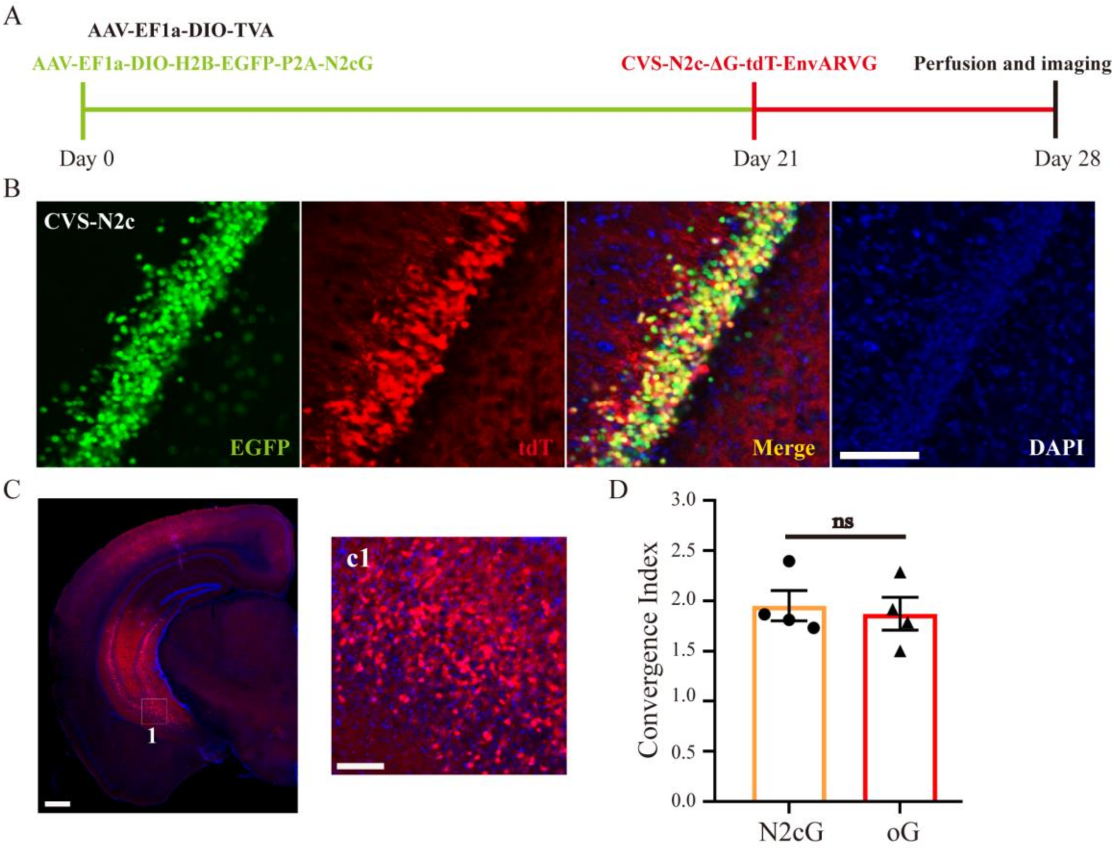


**Fig. S2 Comparison of transsynaptic spread efficiency between CVS-N2c-ΔG/oG and CVS-N2c-ΔG/N2cG.** (A) Schematic diagram of virus injection for trans-mono-synaptic tracing. The adeno-associated viruses carrying Cre-dependent N2cG and TVA were injected into the ventral hippocampal region (vHPC) of Thy1-CRE transgenic mice. After 3 weeks, EnvARVG pseudotyped CVS-N2c-ΔG was injected at the same site respectively. After 1 week, brain slices were processed and imaged by slide scanner. (B) Starter cells at injection site. The green fluorescence signals of N2cG could be co-labeled with the red fluorescence signals of RV. (C) Monosynaptic input neurons in contralateral vHPC labeled by N2cG mediated CVS-N2c-ΔG spread. c1 is a partial enlarged view of figure C. (D) Convergence indices for long-distance input in contralateral vHPC. Statistical values are indicated as mean ± SEM. Significant differences are expressed by the p value. *$P<0.05$, **$P<0.01$, ***$P<0.001$, ns, no significant difference. Scale bars: 100 μm for figure B/c1; 500 μm for figure C.

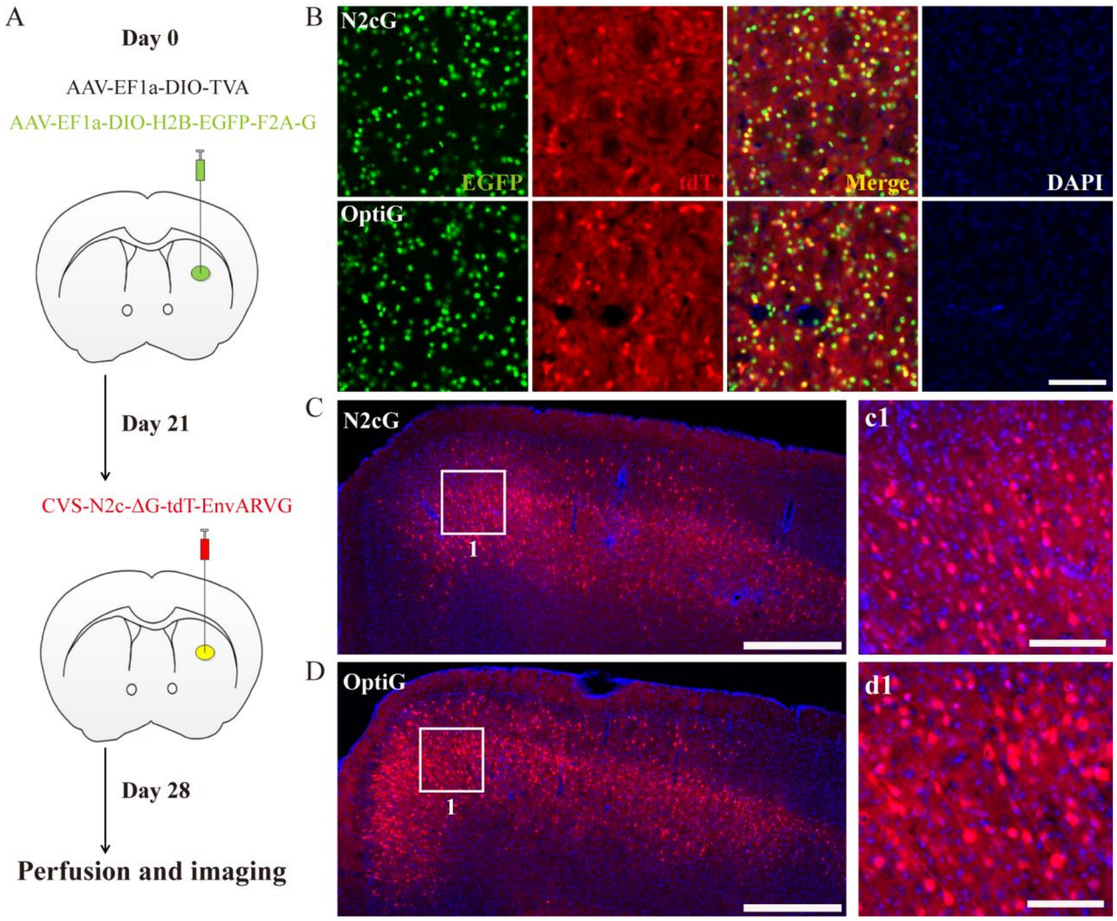


**Fig. S3 Effective transsynaptic spread with optiG mediated CVS-N2c-ΔG.** (A) Schematic diagram of virus injection for trans-mono-synaptic tracing. The adeno-associated viruses carrying Cre-dependent RVG and TVA were injected into the CPu of D2R-Cre transgenic mice. After 3 weeks, EnvARVG pseudotyped CVS-N2c-ΔG was injected at the same site respectively. After 1 week, brain slices were processed and imaged by slide scanner. (B) Starter cells at injection site. The green fluorescence signals of RVG could be co-labeled with the red fluorescence signals of RV. (C) Monosynaptic input neurons in cortex labeled by N2cG mediated CVS-N2C-ΔG spread. c1 is a partial enlarged view of figure C. (D) Monosynaptic input neurons in cortex labeled by optiG mediated CVS-N2C-ΔG spread. d1 is a partial enlarged view of figure D. Scale bars: 100 μm for figure B/c1/d1; 500 μm for figure C/D.

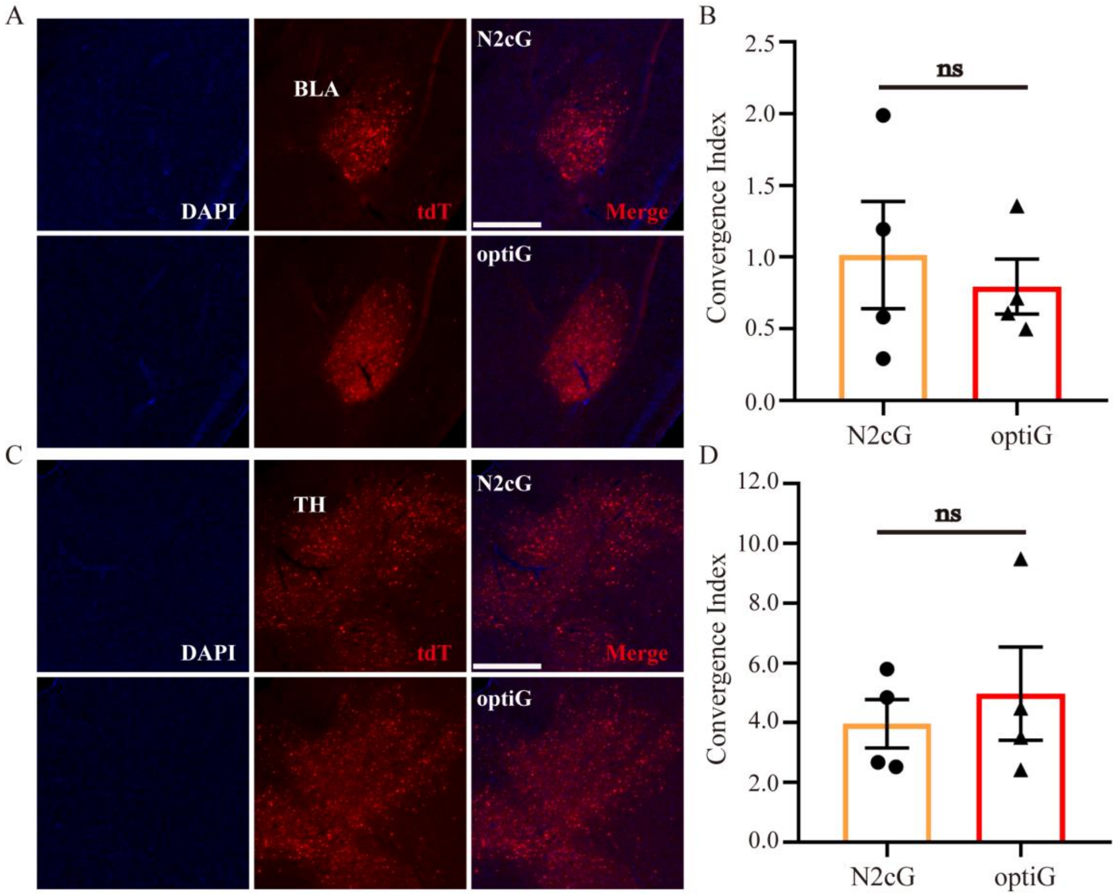


**Fig. S4 Comparison of transsynaptic spread efficiency between CVS-N2c-ΔG/optiG and CVS-N2c-ΔG/N2cG.** (A and C) Coronal sections showing tdTomato$^{+}$ monosynaptic input neurons in BLA (A) and TH (C) to D2R neurons in CPu when using N2cG (top) and optiG (bottom). (B and D) Convergence indices for long-distance inputs in BLA (B) and TH (D) when using N2cG and optiG. Statistical values are indicated as mean ± SEM. Significant differences are expressed by the p value. *P<0.05, **P<0.01, ***P<0.001, ns, no significant difference. Scale bars: 500 μm.
